# Choice activity stabilizes sensory representations and mediates sensorimotor associations in parietal cortex

**DOI:** 10.1101/2020.05.18.101360

**Authors:** Ting-Yu Chang, Raymond Doudlah, Byounghoon Kim, Adhira Sunkara, Meghan Lowe, Ari Rosenberg

## Abstract

Selecting actions which achieve desired goals often requires three-dimensional (3D) representations of the environment. Because the sensory epithelia cannot directly encode the world’s 3D spatial features, sensory signals must be converted into 3D representations. Here we investigated the relationships between the quality of 3D visual representations, choice-related activity, and motor-related activity in the parietal cortex of macaque monkeys using an eight-alternative 3D orientation discrimination task, visually guided saccade task, and laminar probe recordings. We found that choice activity was preferentially carried by caudal intraparietal area neurons with more robust 3D representations. Choice activity further stabilized the 3D representations, rather than attenuating information not directly relevant to the behavioral task (nuisance variables). An experience-dependent, sensorimotor association additionally aligned sensory and saccade direction preferences, particularly for neurons with choice activity. These findings reveal novel roles for choice activity in improving the fidelity of ecologically relevant object representations and mediating sensorimotor associations.

## Introduction

Interactions with the environment require that sensory information be mapped to motor responses. The parietal cortex is an important site of sensorimotor transformations (Rushworth et al., 1997; Buneo et al., 2002; Brovelli et al., 2004; Buneo and Andersen, 2006). Parietal lesions can result in deficits associated with the impaired use of sensory information to create and execute motor plans, as opposed to deficits in sensory processing or action (Pause and Freund, 1989; Pause et al., 1989). They can also produce 3D visual processing deficits (Holmes, 1918; Holmes and Horrax, 1919). Although sensorimotor transformations are often studied using two-dimensional (2D) stimulus paradigms, mappings between 3D object information and motor responses are essential in the natural world. Thus, parietal cortex may have a fundamental role in creating robust 3D representations and mapping them to specific actions.

Within parietal cortex, an important site of 3D sensory processing is the caudal intraparietal (CIP) area. Neurons in CIP are tuned for 3D orientation (Taira et al., 2000; Rosenberg et al., 2013) signaled by multiple cues (Tsutsui et al., 2001; Tsutsui et al., 2002; Rosenberg and Angelaki, 2014b), and perform multisensory processing to achieve gravity-centered object representations (Rosenberg and Angelaki, 2014a). In addition to carrying high-level sensory representations, CIP activity correlates with the short-term memory and perceptual matching of 3D features (Tsutsui et al., 2003), and choices made during a binary orientation discrimination task (Elmore et al., 2019). Furthermore, inactivating CIP impairs 3D feature discrimination (Tsutsui et al., 2001; Van Dromme et al., 2016).

The anatomical projections and effective connectivity of CIP suggest that it may also contribute to goal-directed sensorimotor transformations (Nakamura et al., 2001; Premereur et al., 2015; Van Dromme et al., 2016; Lanzilotto et al., 2019). In particular, CIP projects to areas involved in motor planning and execution, including the lateral intraparietal area (LIP) (Andersen et al., 1992; Bennur and Gold, 2011; Shushruth et al., 2018), anterior intraparietal area (Murata et al., 2000; Baumann et al., 2009; Pani et al., 2014), and V6A (Fattori et al., 2010; Fattori et al., 2012; Breveglieri et al., 2016). However, it is untested if CIP carries motor-related signals, or if its output only contains 3D sensory information.

Here we investigated the relationships between sensory representations, choice-related activity, and motor-related activity in CIP. Neuronal activity was recorded while macaque monkeys performed an eight-alternative forced choice (8AFC) tilt discrimination task with planar surfaces presented at different slants and distances (Chang et al., 2020). The neurons differed in the extent to which their 3D orientation selectivity depended on distance. Choice activity was preferentially carried by neurons with more robust 3D representations, and further stabilized 3D selectivity. Choice tuning was parametric, and the choice and tilt preferences aligned. Motor-related activity was assessed using a visually guided saccade task (Munoz and Wurtz, 1995; Hanes and Schall, 1996). We found that many CIP neurons had saccade direction tuning. Moreover, the surface tilt and saccade direction preferences were mapped onto one another, reflecting a sensorimotor association that was strongest for neurons with choice activity.

These results identify systematic relationships between sensory representations, choice-related activity, and motor-related activity. Neurons that robustly represented the task-relevant information were most strongly coupled with the choices. While choice activity is widely associated with decision processes, the current results reveal novel roles in stabilizing sensory representations and mediating sensorimotor associations. The coupling of sensory and motor functions by single neurons may facilitate sensorimotor transformations over short timescales that promote successful interactions with the environment.

## Results

We tested three hypotheses about the relationships between sensory representations, choice-related activity, and motor-related activity. First, neurons with more robust 3D representations would preferentially carry choice activity. Second, choice activity would stabilize 3D selectivity, as opposed to attenuating information that was not directly relevant to the task at hand (i.e., nuisance variables). Third, sensorimotor associations would be mediated by choice activity.

## Quantifying 3D tilt sensitivity

Testing our hypotheses required a 3D visual discrimination task that could also be used to quantify the robustness of neuronal 3D representations. To this end, we trained two monkeys to perform an 8AFC orientation discrimination task under different viewing conditions that determined the task difficulty (Chang et al., 2020). The task required the monkeys to report which side of a plane was nearest (i.e., the plane’s tilt) through a saccade to one of eight choice targets (**Figure 1A**).

**Figure 1.**
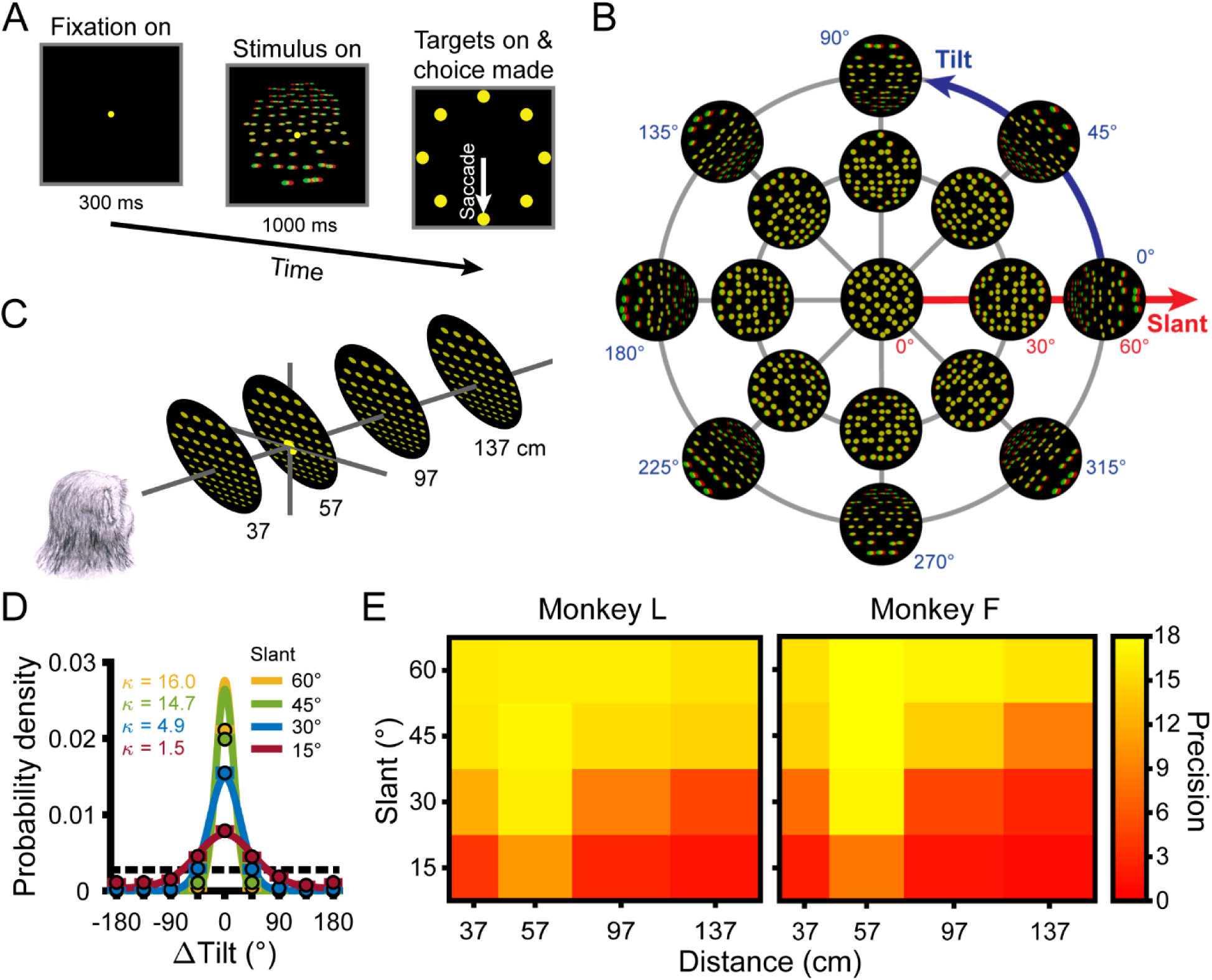
Task, stimuli, and performance. (**A**) Tilt discrimination task. After fixating a target at the center of the screen for 300 ms, a plane was shown for 1 s while fixation was maintained. The fixation target and plane then disappeared and eight choice targets appeared. The plane’s tilt was reported via a saccade to the corresponding target (e.g., the bottom target for a bottom-near plane). (**B**) Tilt and slant. Planes were rendered as random dot stereograms with perspective and stereoscopic cues. A subset of planes are shown here as red–green anaglyphs. For clarity, the number of dots is reduced and the dot size is increased compared to the actual stimuli. (**C**) The planes subtended 20° of visual angle, and were presented at the center of the screen at four distances. The fixation point (depicted here as a larger yellow dot) was always located at 57 cm. (**D**) Error distributions of reported tilts for Monkey L at each slant (distance = 137 cm), calculated over all tilts. Data points show the mean probability of a given ΔTilt (correct choice: ΔTilt = 0°) and SEM (error bars are obscured by the data points) across sessions. Solid curves are von Mises probability density function fits. Taller and narrower densities indicate greater sensitivity. Sensitivities (*κ*, the von Mises concentration parameter) are indicated in the inset. At higher sensitivities, deviation between the data point at ΔTilt = 0° and the density function reflect discrete versus continuous representations of the area between sampled tilts (Chang et al., 2020). Chance performance is marked by the dashed horizontal line. (**E**) Heat maps show mean tilt sensitivity across sessions as a function of slant and distance for each monkey. Yellow hues indicate greater sensitivity.

Tilt and slant are polar coordinates describing 3D surface orientation (**Figure 1B**). Tilt is the angular variable (0° ≤ T < 360°) and specifies the direction that the plane is oriented in depth. For example, T = 0° indicates right-near and T = 90° indicates top-near. Planes were presented at 8 tilts (0° to 315°, 45° steps), corresponding to the 8 choice options. Slant is the radial variable (0° ≤ S ≤ 90°) and specifies the amount of depth variation. There is no depth variation at S = 0°, so tilt is undefined and there is no correct task response. Larger slants indicate greater depth variation. Planes were presented at 5 slants (0° to 60°, 15° steps). Each of the 33 orientations was presented at 4 distances: 37, 57, 97, and 137 cm (**Figure 1C**). Thus, 132 unique surface poses were shown.

Behavioral performance was quantified at each combination of tilt, slant, and distance using the distribution of reported tilt errors (ΔTilt = reported tilt – presented tilt) each session (Monkey L: N = 26 sessions; Monkey F: N = 27) (Chang et al., 2020). Sensitivity was defined as the concentration parameter (*κ*) of the von Mises probability density function fit to the distribution of reported tilt errors (equation 1). For both monkeys, we found that sensitivity significantly depended on slant (generalized linear regression; both *p*-values ≤ 2.9×10^−28^) and distance (both *p*-values ≤ 2.7×10^−5^), but not tilt (linearized into cosine and sine components; all four *p*-values ≥ 0.19) (Fisher, 1995). We therefore summarized sensitivity as a function of slant and distance calculated over all tilts (**Figure 1D, Figure 1―figure supplement 1**). Consistent with our previous findings in which behavioral performance in the 8AFC tilt discrimination task was extensively analyzed over a wide range of 3D poses as well as multiple visual cue conditions (Chang et al., 2020), we found that sensitivity decreased with distance from fixation (57 cm) and increased with slant (**Figure 1E**).

**Figure 1―figure supplement 1.**
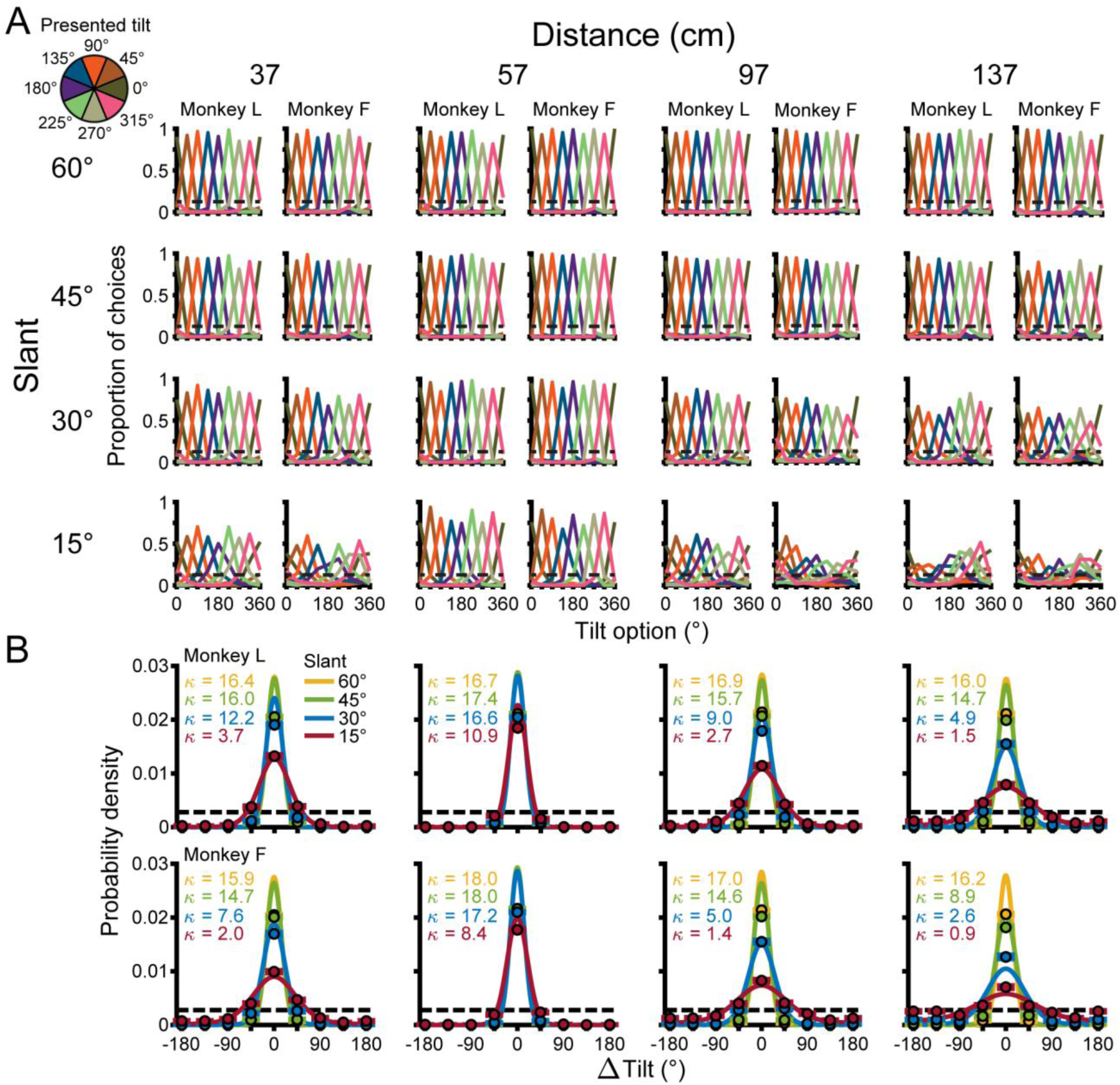
Behavioral performance. (**A**) Proportion of choices made for each tilt option at each combination of tilt (colors), slant (rows), and distance (supercolumns), and each monkey (subcolumns). Chance performance is marked by dashed horizontal lines. Performance was near perfect at combinations of large slants and distances near fixation (57 cm), but approached chance level at small slants and distances far from fixation. Sensitivity was not significantly dependent on tilt (see text for details). (**B**) Error distributions of reported tilts at each combination of slant (colors) and distance (columns), calculated over all tilts (Monkey L: top row; Monkey F: bottom row). Data points show the mean probability of a given ΔTilt (correct choice: ΔTilt = 0°) and SEM (error bars are obscured by the data points) across sessions. Solid curves are von Mises probability densities. Sensitivities are indicated in the insets. Chance performance is marked by dashed horizontal lines.

### CIP responses to 3D surface pose predict tilt sensitivity

We next assessed if the sensory representations in CIP accounted for the monkeys’ tilt sensitivity. While the monkeys performed the task, we recorded from 437 neurons (Monkey L: N = 218; Monkey F: N = 219) using laminar probes (**Figure 2**). Because CIP neurons can carry sensory and choice-related signals (Elmore et al., 2019), we divided the responses into two time windows. The “sensory only” (SO) window started at the median visual response latency (52 ms) and ended at the onset of choice activity (202 ms; calculated below). The “sensory plus choice” (SPC) window, which can include both sensory and choice activity, started at the onset of choice activity and ended at the stimulus offset (1 s). We first analyze responses in the SO window.

**Figure 2.**
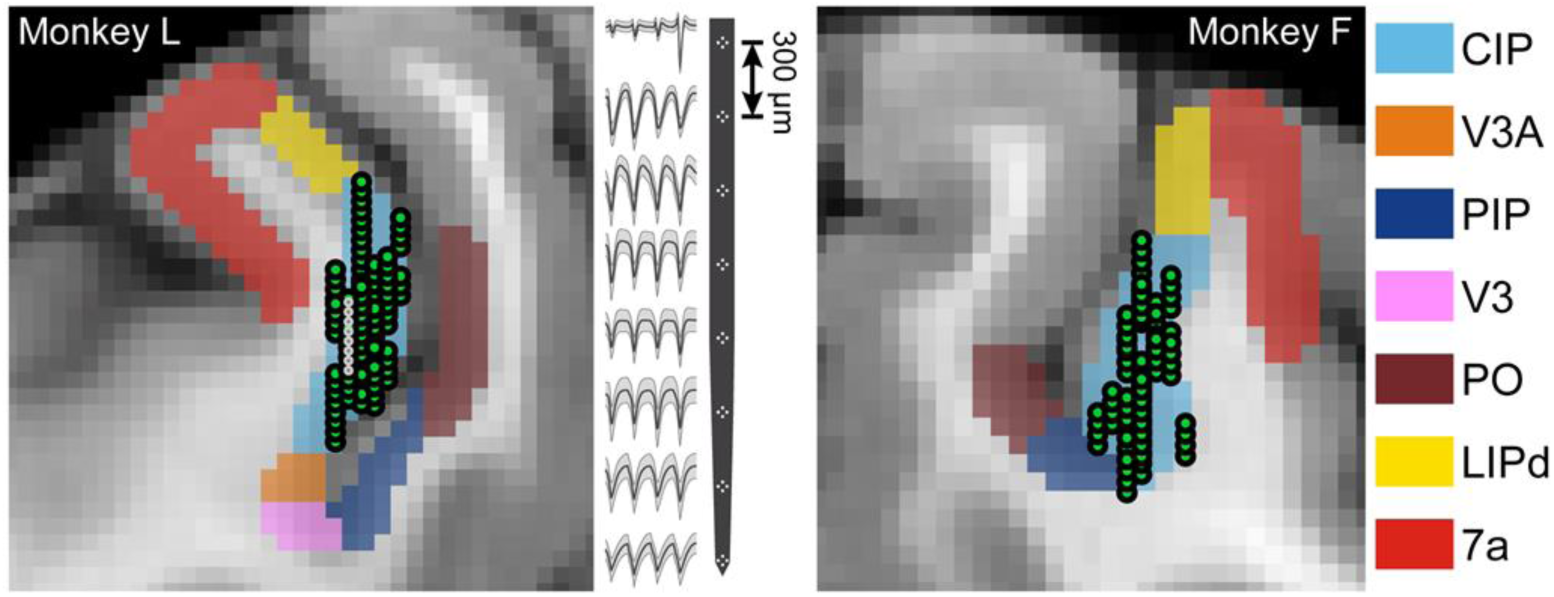
Neuronal recordings. Coronal MRI sections showing the estimated boundaries of CIP (light blue) and neighboring regions (see **Materials and methods**). All recording locations (green circles) are shown, projected along the anterior-posterior (AP) axis onto a single section (Monkey L: AP = −7 mm; Monkey F: AP = −5.5 mm). Spike waveforms from an eight-tetrode recording (locations marked by smaller white circles in the MRI) are shown for Monkey L.

Distinguishing 3D representations from sensitivity to lower-level features (e.g., local binocular disparity cues) that co-vary with changes in object pose is a longstanding problem (Janssen et al., 2000; Nguyenkim and DeAngelis, 2003; Alizadeh et al., 2018; Elmore et al., 2019). These possibilities can be disentangled by assessing how orientation selectivity is affected by the object’s distance when the fixation distance is held constant. A 3D representation is indicated if the orientation selectivity (tuning curve shape) is invariant to the distance (though the gain can change). In contrast, sensitivity to lower-level features is implied if the selectivity changes drastically. The sensory responses of five CIP neurons that portray the range of sensory, choice-related, and motor-related properties we found are shown in **Figure 3**.

**Figure 3.**
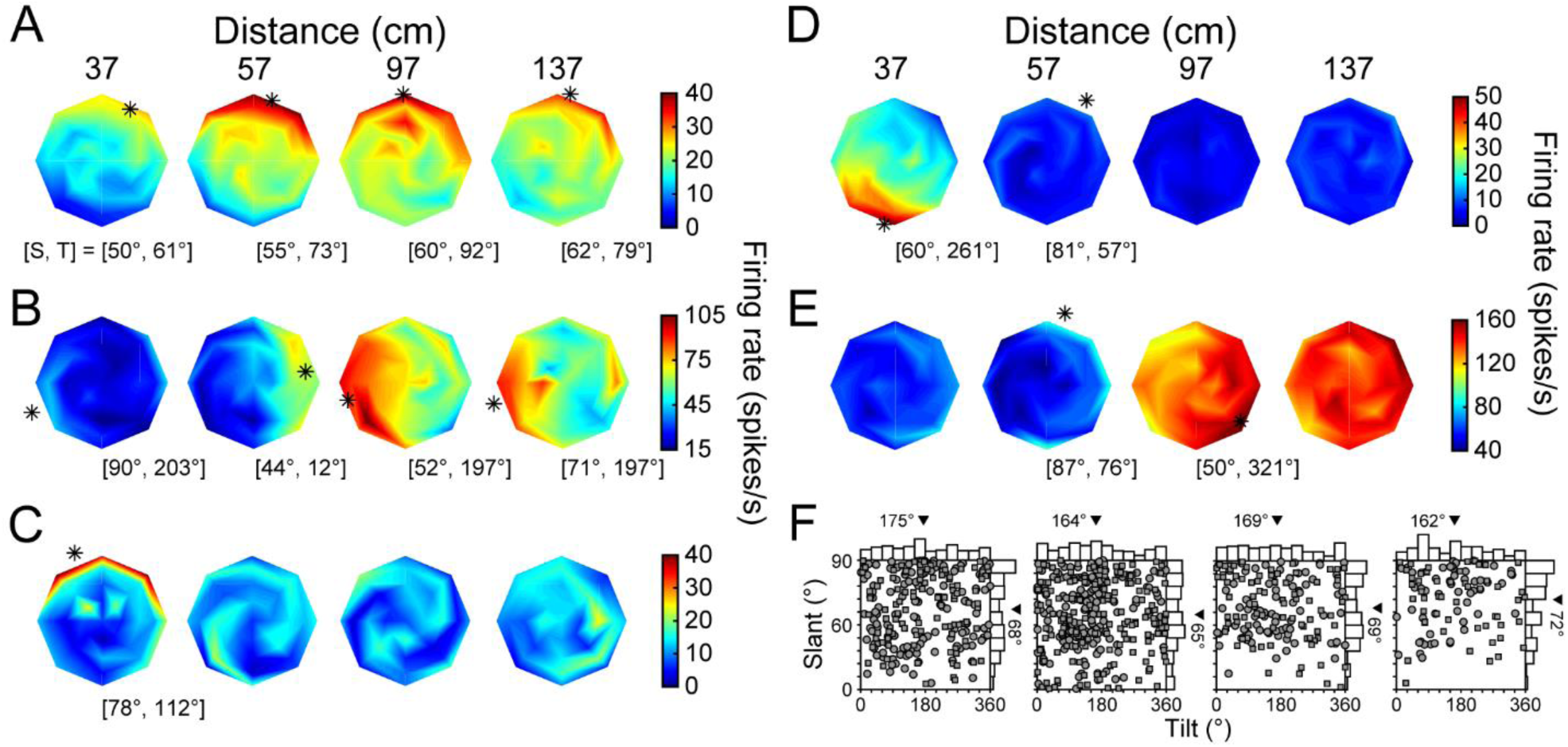
3D orientation tuning at each distance before the onset of choice activity. (**A-E**) Five example neurons. Heat maps show firing rate as a function of tilt (T) and slant (S), plotted using the coordinates illustrated in **Figure 1B**. Red hues indicate higher firing rates. Asterisks mark preferred orientations from Bingham fits (tuned cases only). Some asterisks are not on the discs because the largest tested slant was 60°. (**A**) High tolerance neuron. (**B**) Intermediate tolerance neuron. (**C**) Low tolerance neuron. (**D**) Low tolerance neuron. (**E**) Distance selective neuron with little orientation-related modulation. (**F**) Distribution of tilt and slant preferences at each distance plotted using an equal area projection (Monkey L: circles; Monkey F: squares). Only tuned neurons at each distance are included: 37 (N = 242), 57 (N = 320), 97 (N = 158), 137 (N = 106) cm. Marginal histograms show the tilt and slant distributions. Triangles mark mean values.

In an idealized 3D representation, the orientation tuning (preference and bandwidth, but not gain) is independent of distance. That is, 3D pose tuning will be separable over orientation and distance. To quantify how well CIP activity conformed to this ideal, we fit each neuron’s responses with a separable model (equation 7). A tolerance index was then defined as the average correlation between the responses and fit at each distance. The index quantifies, on a continuum, if a neuron is more selective for 3D pose (values closer to 1) or lower-level visual features (values closer to 0). Some neurons were highly tolerant, indicating robust 3D tuning (**Figure 3A**; Tolerance = 0.88). Other neurons had modest tolerances, suggesting intermediate representations between 3D tuning and lower-level feature selectivity (**Figure 3B**; Tolerance = 0.53). Still others had low tolerances, consistent with lower-level feature selectivity (**Figure 3C,D**; Tolerance = 0.34 and 0.24, respectively). A minority had stronger distance selectivity than orientation selectivity (**Figure 3E**). Across the population, 62 neurons (14%) were distance selective (two-way ANOVA, *p* < 0.05) but not orientation selective (*p* ≥ 0.05).

Vergence eye movements are a potential factor that can influence neuronal responses. To evaluate if our measurements of 3D pose tuning were affected by vergence, we performed an analysis of covariance on the responses of each neuron with tilt (linearized into cosine and sine components), slant, and distance as independent factors and the mean vergence over the analyzed time window as a covariate (DeAngelis and Uka, 2003). Both the SO and SPC windows were included in this analysis because the results were similar for the individual time windows. For only 40 neurons (9%) was there a statistically significant effect of vergence (*p* < 0.05). Moreover, the significance of the tilt, slant, and distance main effects was unchanged for all but 11 neurons (3%) when vergence was included as a covariate. These effects are smaller than those previously reported for disparity tuning in the middle temporal area (DeAngelis and Uka, 2003), and indicate that vergence had a minimal impact on the 3D pose tuning.

We next evaluated if the distribution of orientation preferences during the SO window was distance dependent. Preferences were estimated by fitting a Bingham function to each significant orientation tuning curve (ANOVA, *p* < 0.05, with Bonferroni-Holm correction for each neuron) (Rosenberg et al., 2013). The distribution of tilt preferences was not significantly different from uniform at any distance (*χ*^*2*^; 37, 57, and 97 cm: *p*-values ≥ 0.43; 137 cm: *p* = 0.03, not significant after Bonferroni-Holm correction). At 57 cm (fixation distance), the distribution of slant preferences was not significantly different from uniform (*p* = 0.13). These findings are consistent with our previous results. However, the distributions of slant preferences were significantly different from uniform at 37, 97, and 137 cm (all *p*-values ≤ 1.2×10^−3^, with Bonferroni-Holm correction). At each of these distances, the mean slant preference was larger than at 57 cm (**Figure 3F**).

To determine why the slant preferences were larger at distances closer and further than fixation, we performed pairwise comparisons of the slant preferences of neurons with orientation tuning at adjacent distances. Preferences at 37 and 57 cm were not significantly different (paired t-test, *p* = 0.19, N = 209). For that subset of neurons, we therefore examined the distribution of preferences at 57 cm and found that it was significantly different from uniform (*χ*^*2*^, *p* = 1.5×10^−2^). In contrast, the distribution of preferences for the 111 neurons that represented 57 but not 37 cm was not different from uniform (*p* = 0.24). Thus, neurons that represented 37 cm tended to prefer larger slants. We further found that the preferences of individual neurons tended to increase from 57 to 97 cm (mean ΔSlant = 5.1°; paired t-test, *p* = 1.1×10^−3^, N = 144), as well as from 97 to 137 cm (mean ΔSlant = 4.3°; *p* = 9.7×10^−3^, N = 87). Thus, there was a slight inseparability in the sensory tuning such that the slant preferences tended to increase with distance behind fixation. These results are consistent with the behavioral data which showed that sensitivity fell off more slowly with distance from fixation at larger compared to smaller slants (**Figure 1E**).

We previously found that behavioral tilt sensitivity could be explained by a neuronal probabilistic population code model of perspective and stereoscopic cue integration (Chang et al., 2020). The model predicted a monotonic relationship between behavioral sensitivity and neuronal response gain. We tested this prediction by comparing the tilt sensitivity at each slant–distance combination to the corresponding gain of the tilt tuning curves averaged over all neurons (**Figure 4**). Consistent with the prediction, tilt sensitivity increased as a function of response gain (Monkey L: Spearman r = 0.95, *p* = 2.2×10^−308^; Monkey F: r = 0.87, *p* = 2.2×10^−308^). The finding that response gain was highly predictive of behavioral sensitivity across performance levels ranging from near-chance to near-perfect (**Figure 1E, Figure 1―figure supplement 1**) suggests that CIP activity may constrain the precision of tilt perception.

**Figure 4.**
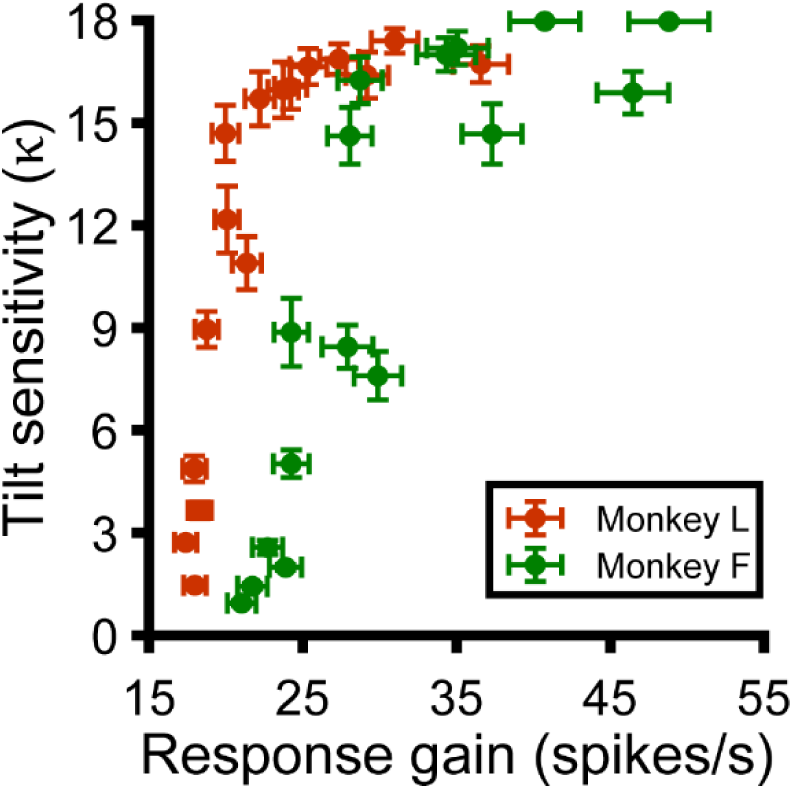
Neuronal response gain predicts behavioral tilt sensitivity. Tilt sensitivity (*κ*) versus response gain for each slant–distance combination (Monkey L: orange; Monkey F: green). Data points show mean and SEM across sessions or neurons. Sensitivity does not exceed *κ* = 18, the upper limit that can be estimated with a 45° tilt sampling interval (Chang et al., 2020).

### Choice activity was parametrically tuned and aligned with the sensory preferences

We next tested each neuron for choice-related activity. An advantage of the 8AFC task was that it could reveal parametric choice tuning, which is not possible with a typical binary choice task. To test for choice-related activity, we only used frontoparallel plane (S = 0°) responses since tilt is undefined at that orientation. To increase the statistical power, we combined responses across all distances after separately z-scoring them at each distance. Responses were then grouped according to the choice that was made on each trial.

We computed eight population-level time courses of choice-related activity, relative to the choice that elicited the maximum response for each neuron (**Figure 5A**). The time course of choice activity had parametric tuning, with an amplitude that symmetrically decreased with greater deviation from the preferred choice (note the similarity of the ±45°, ±90°, and ±135° curves). The onset of choice activity was defined as the first time point after stimulus onset that the curves significantly diverged (ANOVA, *p* < 0.05). All choice analyses were performed using the SPC window, defined from the onset of choice activity (202 ms) to the end of the stimulus presentation (1 s). Results were similar using a shorter window (850 ms – 1 s) that matched the duration of the SO window. Across the population, 201 neurons (46%) had choice activity (ANOVA, *p* < 0.05).

**Figure 5.**
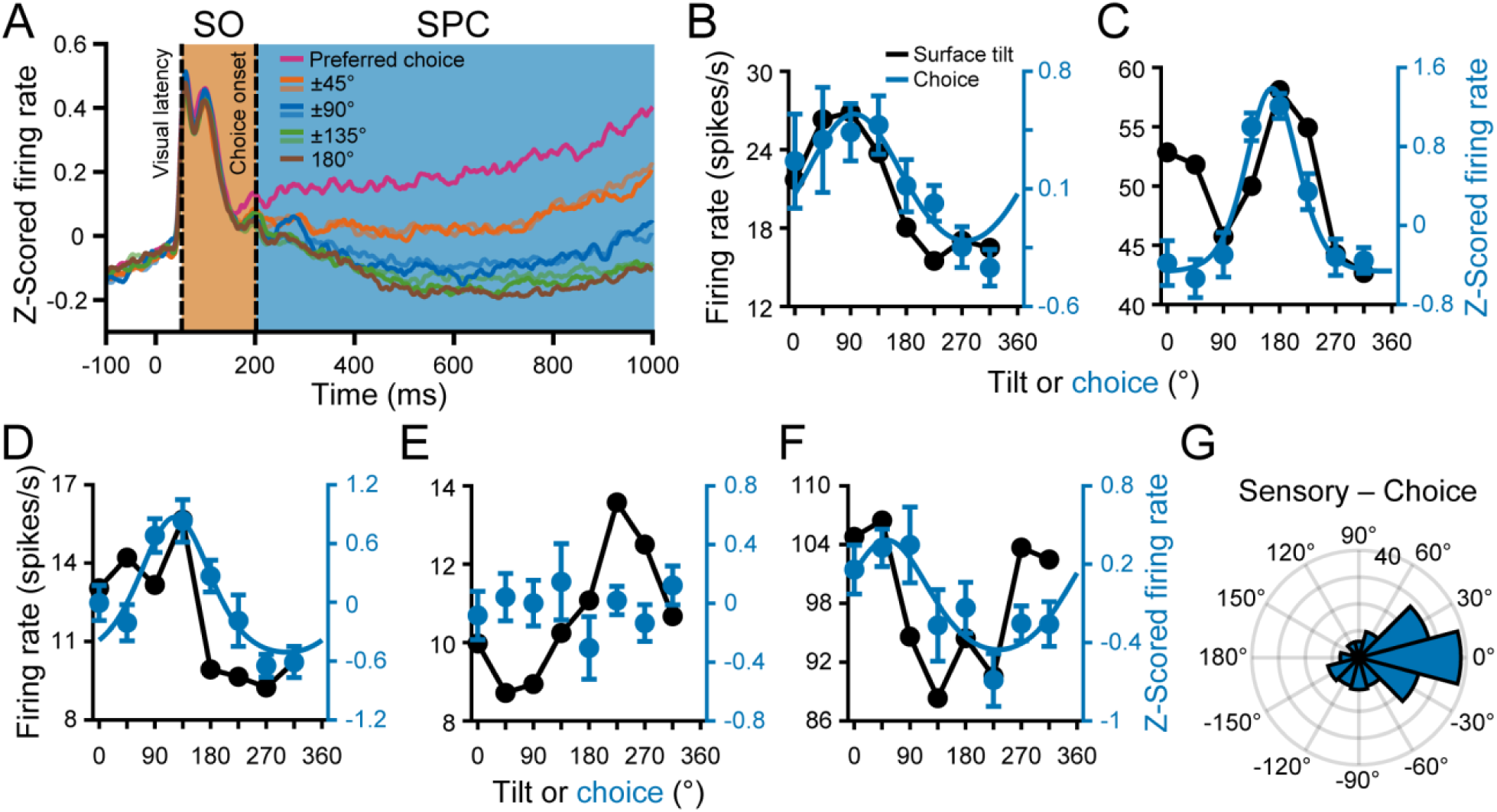
Choice tuning was parametric and aligned with the sensory preferences. (**A**) Time course of choice-related activity. Curves are average z-scored responses for each of the eight choices relative to the preferred choice. Stimulus onset = 0 ms. Vertical dashed lines mark the response latency (52 ms) and onset of choice activity (202 ms). Shaded regions mark the SO (orange) and SPC (blue) windows. (**B-F**) Comparison of surface tilt and choice tuning for the example neurons (**Figure 3A-E**, same order). Black points are surface tilt responses marginalized over slant and distance (SO window). In **C**, the tilt tuning curve has two peaks because the orientation preference changed with distance. Blue points are z-scored choice responses, and the curves are von Mises fits (tuned cases only). (**G**) Difference between principal surface tilt (SO window) and choice preferences (N = 166). The peak at 0° indicates that sensory and choice preferences tended to align.

Choice tuning curves with von Mises fits are shown for the example neurons in **Figure 5B-F** (blue curves). The high (**Figures 3A, 5B**), intermediate (**Figures 3B, 5C**), and first low (**Figures 3C, 5D**) tolerance neurons all had choice activity. The second low tolerance neuron did not have choice activity (**Figures 3D, 5E**). The neuron with stronger distance than orientation tuning also had choice activity (**Figures 3E, 5F**). Across the population, the choice tuning curves were well described by von Mises functions (mean r = 0.88 ± 0.10 SD, N = 201). The mean concentration was *κ* = 3.52 ± 4.74 SD, and the mean half-width at half-height was 52° ± 21° SD.

For comparison, surface tilt tuning curves marginalized over slant and distance (SO window) are shown in **Figure 5B-F** (black curves). The tilt and choice preferences of the example neurons aligned, even for the neuron with stronger distance than orientation tuning. To quantify this relationship across the population, we used the orientation preferences measured at each distance to compute a principal orientation preference for each neuron (SO window; see **Materials and methods**). We then compared the principal surface tilt and choice preferences from the von Mises fits (**Figure 5G**). The median circular difference between the preferences was −8.3° and not significantly different from 0° (circular median test, *p* = 0.24; N = 166 neurons with orientation and choice tuning), indicating that sensory and choice preferences tended to align.

### Changes in 3D selectivity associated with choice activity

We next tested if choice activity was associated with changes in 3D selectivity. To start, we calculated the correlation between tuning curves in the SO and SPC windows (over all 132 poses). The mean correlation was r = 0.49 ± 0.27 SD (N = 437), suggesting that tuning was similar in the two time windows but not identical. We hypothesized that the changes reflected a stabilizing effect of choice activity on 3D tuning, but also considered that choice might attenuate selectivity for slant and distance (nuisance variables during tilt discrimination), leaving only selectivity for the task-relevant variable (tilt) intact.

To illustrate how selectivity changed after the onset of choice activity, the orientation tuning at each distance is shown for the example neurons during the SPC window in **Figure 6A-E**. First consider the three neurons with strong orientation and choice tuning. The selectivity of the neuron with high tolerance in the SO window changed little (**Figures 3A, 6A**), and its tolerance slightly increased: ΔTolerance = 0.08. Strikingly, the orientation tuning of the neuron with intermediate tolerance in the SO window aligned across distances in the SPC window, matching the tuning at the preferred distance (**Figures 3B, 6B**), and its tolerance greatly increased: ΔTolerance = 0.35. For the neuron with low tolerance in the SO window, responses at non-preferred distances remained relatively weak in the SPC window. However, the orientation tuning was now significant at each distance and the preferences were similar (**Figures 3C, 6C**), resulting in a large increase in tolerance: ΔTolerance = 0.46. Next consider the neuron with low tolerance during the SO window and no choice activity. The selectivity of this neuron changed relatively little during the SPC window (**Figures 3D, 6D**). Lastly, the distance selective neuron continued to respond most strongly to the furthest distances (**Figures 3E, 6E**). During the SPC window, 20 neurons (5%) were distance selective (two-way ANOVA, *p* < 0.05) but not orientation selective (*p* ≥ 0.05), compared to 62 (14%) in the SO window.

**Figure 6.**
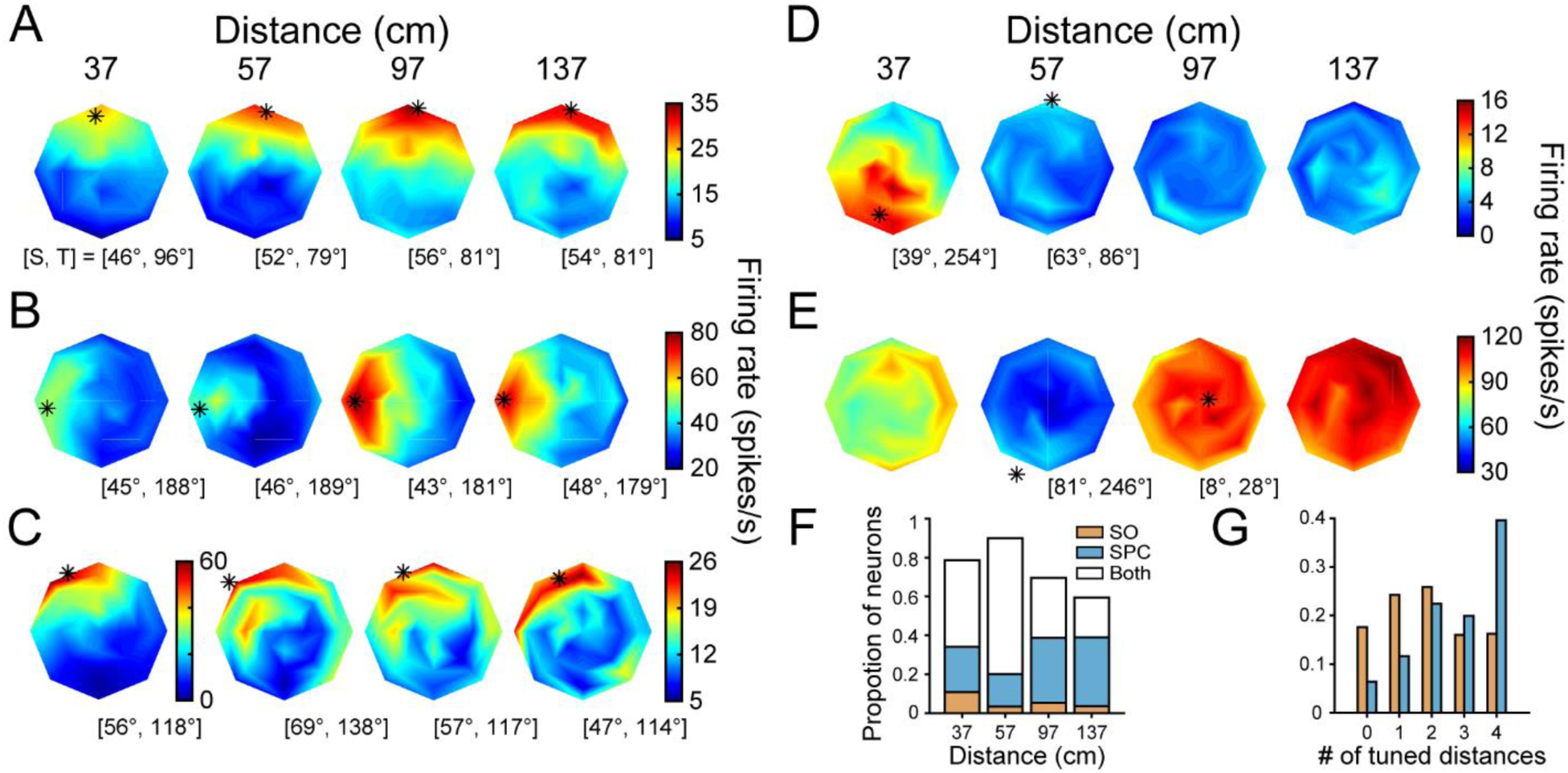
3D orientation tuning at each distance after the onset of choice activity. (**A-E**) Responses of the example neurons (**Figures 3A-E, 5B-F**, same order). (**A**) Tolerance increased from 0.88 (SO window) to 0.96 (SPC window). (**B**) Tolerance increased from 0.53 to 0.88. Note the changes in selectivity at 37 and 57 cm. (**C**) Tolerance increased from 0.34 to 0.80. The number of distances with significant orientation tuning increased from one to four. (**D**) Tolerance increased from 0.24 to 0.33. (**E**) Distance selective neuron. (**F**) Proportion of neurons with orientation tuning at each distance during the SO window only (orange), SPC window only (blue), or both (white). The proportion of neurons with tuning decreased with distance from fixation (57 cm), consistent with the pattern of behavioral sensitivity (**Figure 1E**). (**G**) Proportion of neurons with orientation tuning at each possible number of distances during the SO (orange) and SPC (blue) windows.

Slant and distance tuning persisted in the SPC window for the example neurons, suggesting that choice activity did not attenuate information that was not directly relevant to the task. This finding was typical across the population. For each neuron and time window, we tested for slant and distance tuning (4-way ANOVA with tilt linearized into cosine and sine components, *p* < 0.05). The number of neurons with slant tuning was 334 (76%) in the SO window and 370 (85%) in the SPC window. The number of neurons with distance tuning was 414 (95%) in the SO window and 412 (94%) in the SPC window. These results are inconsistent with choice activity attenuating the representation of nuisance variables since the prevalence of slant and distance tuning did not decrease during the SPC window. Instead, choice activity was associated with an increase in the number of neurons with significant orientation tuning (ANOVA, *p* < 0.05) at each distance (**Figure 6F**). Indeed, individual neurons often had orientation tuning at two distances during the SO window but all four distances during the SPC window (**Figure 6G**).

### Choice activity was carried by more tolerant neurons and stabilized 3D selectivity

We next tested if neurons with robust 3D tuning preferentially carried choice activity. To do so, we compared the SO window tolerances of neurons with and without choice activity (**Figure 7A**). The mean tolerance was greater for neurons with choice activity (0.61) than without (0.56). Although the difference was not large, it was significant (ANOVA followed by Tukey’s HSD test, *p* = 0.02), indicating that choice activity was preferentially carried by neurons with more robust 3D tuning. We then tested if choice activity further stabilized 3D selectivity by assessing if tolerance increased in the SPC window. The difference in mean tolerance between neurons with (0.72) and without (0.60) choice activity increased in the SPC window (**Figure 7B**) and was significant (*p* = 3.8×10^−9^). The greater difference was due to an increase in tolerance between time windows for neurons with choice activity (*p* = 3.8×10^−9^; **Figure 7C**). The tolerance of neurons without choice activity did not significantly change (*p* = 0.06). We additionally found that the SPC window tolerance of neurons with choice activity was correlated with the strength of choice activity (log of the tuning curve gain): r = 0.45, *p* = 2.8×10^−11^.

**Figure 7.**
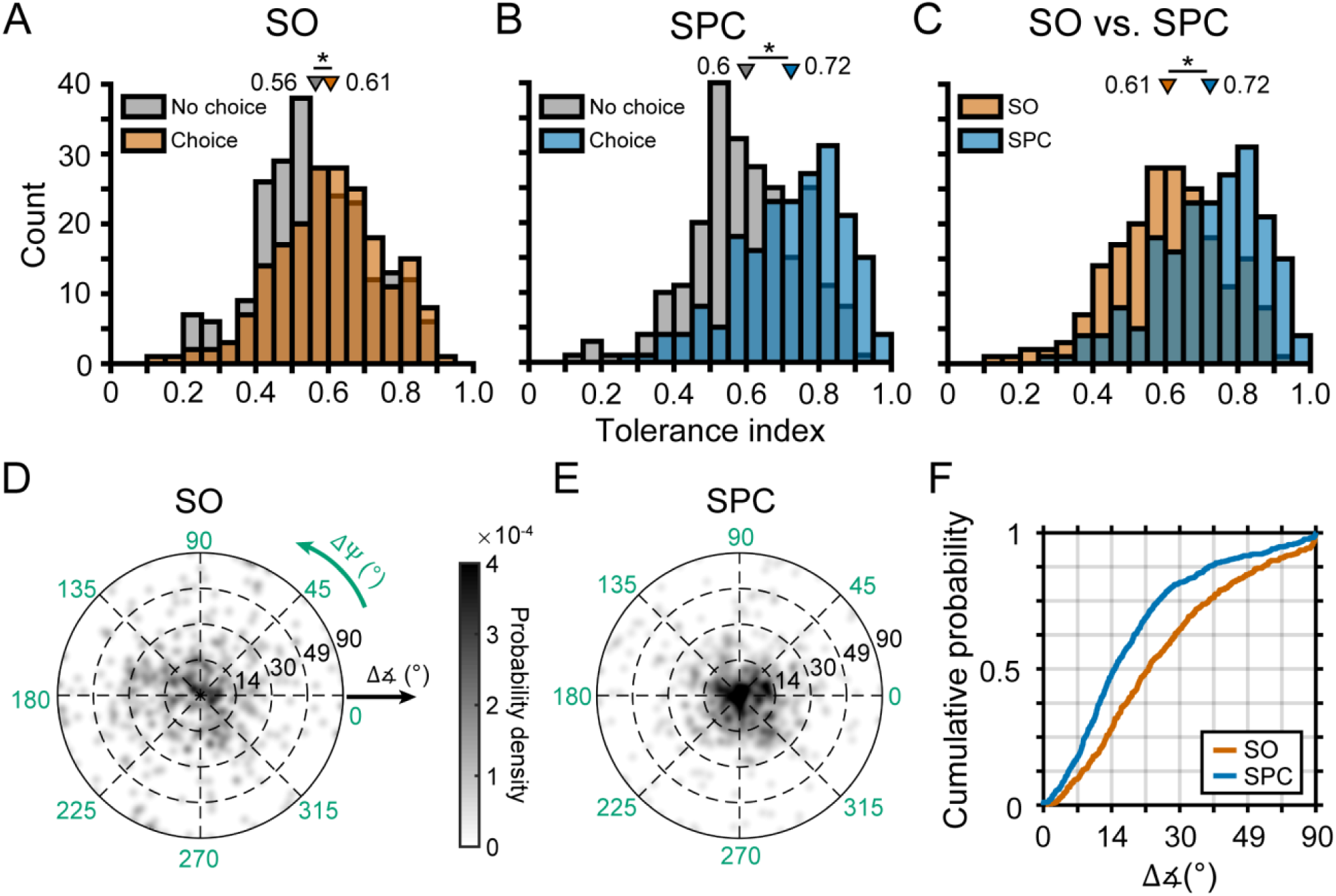
Choice activity was carried by more tolerant neurons and stabilized 3D selectivity. (**A**) Comparison of tolerance in the SO window for neurons with (orange) and without (gray) choice activity. (**B**) Comparison of tolerance in the SPC window for neurons with (blue) and without choice activity. (**C**) Comparison of tolerance between the SO and SPC windows for neurons with choice activity. In **A-C**, triangles mark mean tolerances. Asterisks indicate significant differences. (**D**) Probability density for the deviations in orientation preference (direction: ΔΨ; angle: Δ∡) from the principal orientation (calculated using both time windows) during the SO window (Gaussian smoothing kernel, σ = 0.025). (**E**) Same as **D** but for the SPC window. (**F**) Cumulative density functions for the angular deviations (radial distance in **D,E**) in the SO and SPC windows. In **C-F**, only neurons with choice activity are included.

To further characterize the effects of choice activity on 3D selectivity, we quantified the differences in orientation preference across distance in each time window. For each neuron with choice activity, we calculated a principal orientation preference using both time windows (see **Materials and methods**). Deviations in orientation preference from the principal orientation are plotted for each time window in **Figure 7D,E**. In these plots, the angular variable is the direction that the preferred orientation at a given distance deviated from the principal orientation (ΔΨ), and the radial variable is the angular deviation (Δ∡). The origin indicates no difference, and points on the outer ring indicate the maximal difference (90°). The deviations were not significantly different from the origin in either time window (test for a specified principal axis; both *p*-values ≥ 0.09) (Fisher et al., 1993). To assess how much the orientation preferences deviated from the principal orientation, we calculated cumulative density functions for the angular deviations (**Figure 7F**). The mean deviation was greater in the SO (28.3°) than the SPC (20.4°) window, and the cumulative densities significantly differed (Kolmogorov-Smirnov test, *p* = 1.4×10^−10^). Thus, the orientation preferences became more similar across distance after the onset of choice activity. We repeated this analysis for neurons without choice activity, and found that the cumulative densities for the two time windows were not significantly different (*p* = 0.91). Thus, 3D selectivity stabilized in the SPC window, but only for neurons with choice activity.

### Sensorimotor associations in CIP are mediated by choice activity

We lastly conducted the first test of motor-related activity in CIP. Anatomical and effective connectivity studies indicate that CIP receives (direct or indirect) input from V3A and projects to LIP (Nakamura et al., 2001; Premereur et al., 2015; Van Dromme et al., 2016), two areas with saccade-related activity (Andersen et al., 1992; Nakamura and Colby, 2000). We therefore tested for saccade direction tuning using a visually guided saccade task (Munoz and Wurtz, 1995; Hanes and Schall, 1996) (**Figure 8A**).

**Figure 8.**
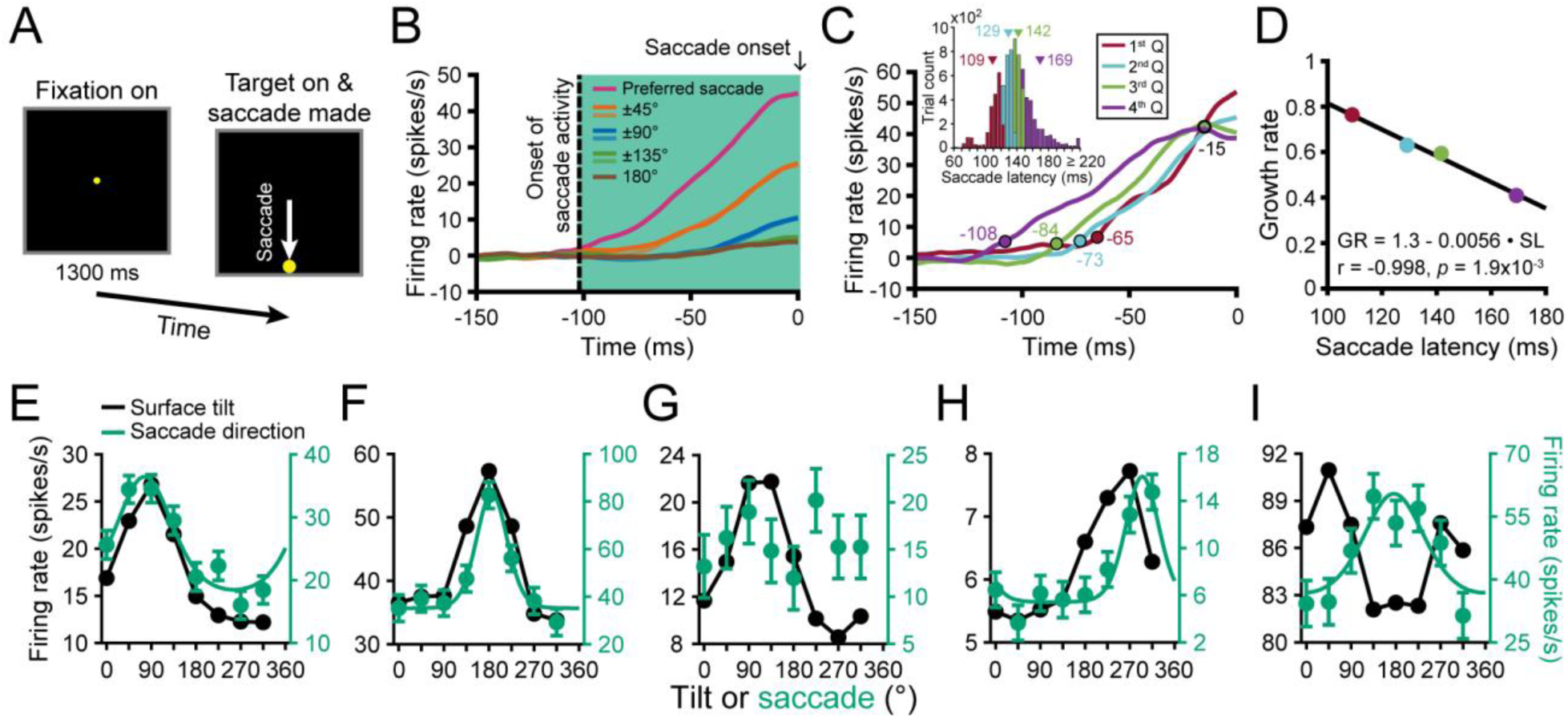
Saccade-related activity and sensorimotor associations. (**A**) Saccade task. After fixating a target at the center of the screen for 1.3 s, the fixation target disappeared and a saccade target appeared at one of eight locations coinciding with the choice targets in the discrimination task. (**B**) Time course of saccade-related activity. Curves are average responses for each of the eight saccade directions relative to the preferred direction. Saccade onset = 0 ms. Vertical dashed line marks the onset of saccade activity (−102 ms). Shaded region marks the window used to assess saccade activity. (**C**) Time course of saccade-related activity depending on the saccade latency in quartiles (Q). Colored circles mark the start of activity. Open black circle marks a putative saccade threshold. The times are indicated next to each circle. Inset shows the histogram of saccade latencies with quartiles colored. Triangles mark mean latencies. (**D**) Inverse linear relationship between the growth rate (GR) of saccade-related activity and mean saccade latency (SL) across quartiles. (**E-I**) Comparison of surface tilt and saccade direction tuning curves for the example neurons (**Figures 3A-E, 5B-F, 6A-E**, same order). Black points are surface tilt responses marginalized over slant and distance (SO+SPC windows). Green points are saccade direction responses, and the curves are von Mises fits (tuned cases only).

We computed eight population-level time courses of saccade-related activity, relative to the saccade direction that elicited the maximum response for each neuron (**Figure 8B**). The time course of saccade-related activity had parametric tuning, with an amplitude that symmetrically decreased with greater deviation from the preferred direction. Significant saccade direction selectivity began 102 ms before saccade onset (ANOVA, *p* < 0.05). We further found that the activity predicted the saccade timing. For each neuron, we labeled every trial in which a saccade was made in the preferred direction according to the saccade latency in quartiles. Time courses were then calculated for each quartile (**Figure 8C**; inset shows the latency histogram). The four curves approximately intersected 15 ms before saccade onset, suggestive of a saccade initiation threshold (∼42 spikes/s, black circle). Moreover, the activity increased more slowly when the saccade latency was longer (colored circles mark when each curve significantly deviated from baseline; ANOVA, *p* < 0.05). For each curve, we computed the growth rate (linear slope) between the start of activity and the putative saccade threshold. Consistent with frontal eye field findings (Hanes and Schall, 1996), there was an inverse linear relationship between the growth rate and mean saccade latency (**Figure 8D**). Thus, saccade-related activity in CIP functionally correlated with both the saccade direction and timing during the visually guided saccade task.

Across the population, 274 neurons (∼63%) had saccade direction tuning (ANOVA, *p* < 0.05). The tuning curves were well described by von Mises functions (mean r = 0.91 ± 0.10 SD). The mean concentration parameter was *κ* = 4.67 ± 4.84 SD, and the mean half-width at half-height was 42° ± 17° SD. Saccade direction tuning curves with von Mises fits are shown for the example neurons in **Figure 8E-I** (green curves). The first two neurons had orientation (**Figures 3A,B, 6A,B**) and choice (**Figure 5B,C**) tuning, as well as saccade direction tuning (**Figure 8E,F**). The third neuron had orientation (**Figures 3C, 6C**) and choice (**Figure 5D**) tuning, but not saccade direction tuning (**Figure 8G**). The fourth neuron, which had orientation (**Figures 3D, 6D**) but not choice (**Figure 5E**) tuning, had saccade direction tuning (**Figure 8H**). Lastly, the neuron with stronger distance than orientation tuning (**Figures 3E, 6E**) had both choice (**Figure 5F**) and saccade direction (**Figure 8I**) tuning, but the choice and saccade preferences were not aligned. These various differences illustrate that choice and saccade response properties were dissociable.

For comparison, surface tilt tuning curves marginalized over slant and distance (SO+SPC windows) are shown in **Figure 8E-I** (black curves). The tilt and saccade direction preferences of the example neurons with strong orientation and choice tuning were well aligned (**Figure 8E,F**), implying a sensorimotor association at the individual neuron level. The preferences were also reasonably well aligned for the neuron with strong orientation but not choice tuning (**Figure 8H**). They were not aligned for the neuron with stronger distance than orientation tuning (**Figure 8I**). To quantify the sensorimotor association, we compared the principal tilt (SO+SPC windows) and saccade direction preferences from the von Mises fits. Across the population, the preferences aligned both for neurons without (**Figure 9A**) and with (**Figure 9B**) choice activity. For neurons without (with) choice activity, the median circular difference between the preferences was −0.9° (−1.6°) and not significantly different from 0° (circular median test, both *p*-values ≥ 0.61). Although the preferences tended to align regardless of choice activity, the distribution was significantly wider for neurons without (circular variance = 0.74) than with (0.36) choice activity (two-sample concentration difference test, *p* = 3.2×10^−5^) (Fisher, 1995), indicating that the sensorimotor association was strongest for neurons with choice activity.

**Figure 9.**
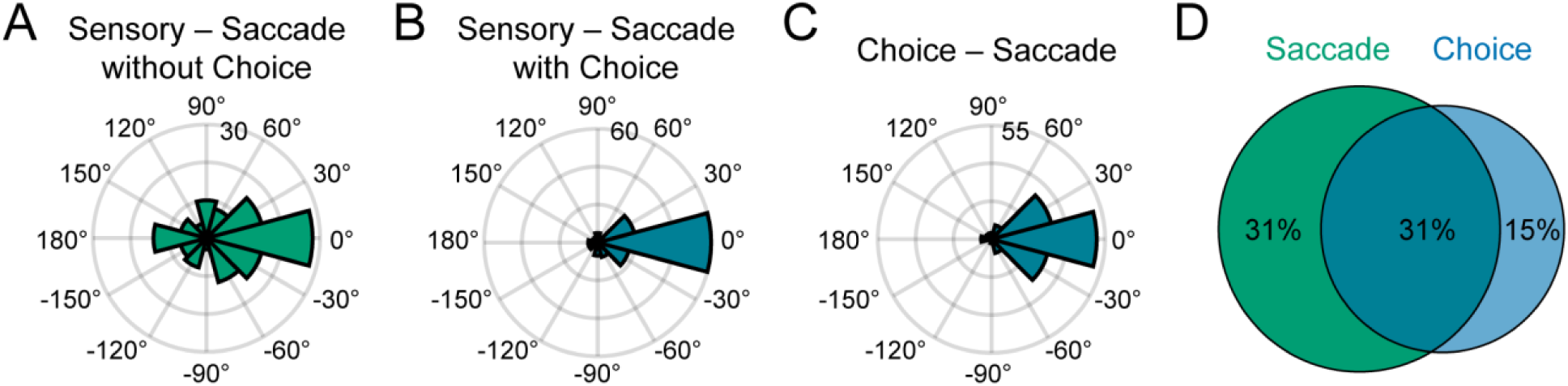
Sensorimotor associations were mediated by choice activity. (**A**) Differences between principal surface tilts and preferred saccade directions for neurons without choice activity (N = 131). (**B**) Same as **A** but for neurons with choice activity (N = 135). (**C**) Differences between choice and saccade direction preferences (N = 137). (**D**) Venn diagram showing the proportion of neurons with saccade activity only (green), choice activity only (blue), and both (teal).

Across the population, 137 neurons (31%) had both choice and saccade direction tuning. The median circular difference between the preferences was 2.9° and not significantly different from 0° (circular median test, *p* = 0.61), indicating that the preferences tended to align (**Figure 9C**). There were also many neurons with saccade activity only (137, 31%) or choice activity only (64, 15%), further indicating that saccade and choice responses were dissociable (**Figure 9D**).

## Discussion

Interacting with the environment requires the creation of robust representations of 3D information that the epithelia cannot directly sense, and mapping those representations to motor responses. We investigated the relationships between the quality of 3D representations, choice-related activity, and motor-related activity in CIP. The behavioral task required the monkeys to report the tilt of a plane, regardless of its slant or distance. As such, slant and distance were nuisance variables that were not of direct interest but modulated task performance. Rather than attenuating selectivity for the nuisance variables, choice activity improved the fidelity of the 3D representation. Thus, the low-dimensional task (report the tilt) did not reduce the dimensionality of the neuronal representation of the multi-dimensional (tilt, slant, and distance) stimulus. This may enable more robust 3D perception and flexible processing. For example, if choice activity attenuated selectivity for features not directly relevant to the task at hand, performance would be impaired if the task and relevant information unexpectedly changed.

The monkeys were trained to report which side of a plane was nearest, whereas a previous CIP study had monkeys report which side was furthest (Elmore et al., 2019). In both studies, surface tilt and choice preferences aligned. For example, given a neuron preferring a bottom-near tilt, the preferred choice report tended to be the lower target in this study but the upper target in the previous study. Here, we further found that many CIP neurons have saccade-related activity, that saccade direction preferences aligned with the sensory and choice preferences, and that choice- and saccade-related activities were dissociable. These findings together reveal a flexible, experience-dependent mapping between sensory, choice-, and motor-related activity, indicating that experience has ongoing effects on the functional properties of CIP neurons, as occurs in downstream area LIP (Freedman and Assad, 2006; Law and Gold, 2008; Bennur and Gold, 2011).

### Building selectivity for 3D object pose

Transforming 2D retinal images into 3D representations is a nonlinear optimization problem (Hartley and Zisserman, 2003). Neurons in CIP achieve 3D selectivity by integrating stereoscopic and perspective cues (Tsutsui et al., 2001; Tsutsui et al., 2002; Rosenberg and Angelaki, 2014b). A stereoscopic representation of 3D orientation that is tolerant to distance requires an encoding of relative disparity gradients that takes into account the nonlinear relationship between depth and disparity. We found that the orientation selectivity of many CIP neurons was highly tolerant to distance. That tolerance cannot simply reflect tuning for perspective cues (even though they were independent of distance) because the vast majority of neurons had significant distance tuning. Our findings thus reveal a high-level representation of 3D object pose created using relative disparity computations. How might this representation be achieved?

Functional properties and connectivity data suggest that tuning for 3D object pose is built hierarchically. Neurons in V1 represent local absolute disparities (Cumming and Parker, 1997, 1999, 2000). A transformation from absolute to relative disparity selectivity likely proceeds in V2 (Thomas et al., 2002) and V3A where absolute disparity signals may still predominate (Anzai et al., 2011) but the representation of disparity gradients may begin (Elmore et al., 2019). The 2D- to-3D transformation likely progresses further in the posterior intraparietal area where selectivity for local relative disparity gradients may arise (Alizadeh et al., 2018), ultimately achieving 3D object pose tuning in CIP.

Recent studies identified brain regions that represent the 3D structure of the environment (Vaziri et al., 2014; Lescroart and Gallant, 2019). How 3D object and environment representations interact at the neuronal level is currently unknown. To characterize sensory and sensorimotor processing under more ecologically realistic contexts, it will be useful to study how CIP represents the pose of more naturalistic objects with which the animals interact, and if that representation is shaped by the surrounding environment.

### Separable orientation and distance tuning

Distinguishing 3D representations from lower-level feature selectivity is a longstanding problem (Janssen et al., 2000; Nguyenkim and DeAngelis, 2003; Alizadeh et al., 2018; Elmore et al., 2019). Showing that 3D orientation selectivity is relatively invariant to distance when the fixation distance is held constant is a common criterion to conclude 3D tuning. This fundamental test was never previously performed in CIP. We found that slant preferences tended to slightly increase with greater distance behind fixation. Although the increase was small, the finding indicates that orientation and distance tuning was not strictly separable. The behavioral data likewise showed that the monkeys’ sensitivity fell off more slowly with distance from fixation at larger compared to smaller slants. This behavioral finding implies that larger slants were required to elicit robust stimulus-selective neuronal responses at distances further from fixation, predicting the observed increase in slant preference and suggesting that strictly separable tuning may not be a reasonable expectation. It is unlikely that this conclusion rests upon using stimuli with fixed slants as opposed to fixed disparity gradients. Although the disparity gradient signaling a given slant is distance dependent (due to the nonlinear relationship between depth and disparity), using fixed disparity gradients would mean that the 3D shape of the stimuli would change with distance. As such, regardless of whether fixed slants or disparity gradients are used, it is unlikely that tuning will be strictly separable over orientation and distance.

### Potential origins of choice-related activity

Choice activity is traditionally associated with feedforward contributions to perception (Celebrini and Newsome, 1994; Britten et al., 1996; Dodd et al., 2001; Nienborg and Cumming, 2006; Gu et al., 2007). This possibility has been questioned on the basis that choice activity, attentional effects, and correlated activity can be conflated (Cumming and Nienborg, 2016). Existing choice activity measures likely reflect combinations of these factors (Haefner et al., 2013; Gu et al., 2014). We found that surface tilt and choice preferences typically aligned. This is expected for a feedforward origin of choice activity, and contrasts with the potentially non-specific effects of a feedback origin, such as proposed for the ventral intraparietal area (Zaidel et al., 2017). We further found that choice activity was preferentially carried by neurons which robustly represented the task-relevant sensory information, and that the strength of choice activity increased with sensory tolerance. The extent to which subcortical vestibular neurons resolve the gravito-inertial acceleration ambiguity necessary to represent translation independent of head tilt (Dakin and Rosenberg, 2018) similarly correlates with the strength of choice activity (Liu et al., 2013). Across modalities, these results reveal that neurons which have resolved fundamental ambiguities about the sensory information that an animal is actively discriminating preferentially carry choice activity. This may reflect a feedforward origin of choice activity, such that neurons with more robust tuning for task-relevant information have greater weight in the neural readout. Indeed, decoding neurons that are more tolerant to nuisance variables may simplify readout since marginalizing out non-relevant features would be more straightforward.

### Sensorimotor associations

We discovered two roles for CIP beyond sensory processing. First, we showed that CIP activity functionally correlates with both the direction and timing of visually guided saccades, implicating the area in visuomotor control for the first time. Second, we found sensorimotor associations at the single neuron level, which were strongest for neurons with choice activity (i.e., those with more robust 3D selectivity). This finding may reflect that neurons with robust sensory representations and stronger sensorimotor associations have greater weight in determining motor responses. Associating robust sensory and motor activity at the single neuron level may help ensure successful and timely interactions with the environment since the same neurons representing the relevant sensory information also signal the appropriate motor response. While our findings implicate CIP in visuomotor functioning, the area is also connected with prehensile areas, suggesting that its sensorimotor functions may be more general. This possibility is consistent with recent findings implicating human parietal regions that integrate visual orientation and saccade signals in the updating of grasp plans during eye movements (Baltaretu et al., 2020).

These findings reveal previously unrecognized roles for choice activity in improving the fidelity of high-level sensory representations and mediating sensorimotor associations, and show that CIP achieves a complete and flexible sensorimotor association for transforming information about the 3D world into overt behavior.

## Materials and methods

### Animal preparation

All surgeries and experimental procedures were approved by the Institutional Animal Care and Use Committee at the University of Wisconsin–Madison, and in accordance with NIH guidelines. Two male rhesus monkeys (*Macaca mulatta*; Monkey L: 6 years of age, ∼9.5 kg; Monkey F: 5 years, ∼6.4 kg) were surgically implanted with a Delrin ring for head restraint and attaching a removable recording grid for guiding electrodes (Rosenberg et al., 2013). After recovery, the monkeys were trained to sit in a primate chair with head restraint, and to fixate visual targets within 2° version and 1° vergence windows. Vergence was enforced during the experiments, which is often not the case in 3D vision studies (e.g., Sanada and DeAngelis, 2014), and was also factored into the analyses using ANCOVAs to test for main effects of stimulus tuning with vergence included as a covariate (DeAngelis and Uka, 2003).

### Experimental control and stimulus presentation

Experimental control was performed using the open-source REC-GUI software (Kim et al., 2019). Stimuli were rendered using Psychtoolbox 3 (MATLAB R2016b; NVIDIA GeForce GTX 970). They were rear-projected onto a polarization preserving screen (Stewart Film Screen, Inc.) using a DLP LED projector (PROPixx; VPixx Technologies, Inc.) with 1,280 × 720 pixel resolution (70° × 43°) at 240 Hz (120 Hz/eye). The screen was positioned 57 cm from the monkey. Polarized glasses were worn. A phototransistor circuit was used to confirm the synchronization of left and right eye images, and to align neuronal responses to the stimulus onset. Eye positions were monitored optically at 1 KHz (EyeLink 1000 plus, SR Research).

### Visual stimuli

The stimuli were the same as the combined-cue stimuli in our previous work (Chang et al., 2020). They consisted of 250 nonoverlapping dots uniformly distributed across a plane. The stimulus envelope was a 20° diameter circle on the screen. The background was gray (11.06 cd/m^2^) and the dots were bright (35.1 cd/m^2^), measured through the glasses (PR-524 LiteMate, Photo Research). Planes were presented at all combinations of eight tilts (0° to 315°, 45° steps), five slants (0° to 60°, 15° steps), and four distances (37, 57, 97, and 137 cm). Fixation was always at screen distance (57 cm). At 37 cm, all dots were in front of the fixation target. At 57 cm, the dots were distributed about the fixation target. At 97 and 137 cm, all dots were behind fixation. Presenting the stimuli in front of, distributed about, and behind fixation prevented the monkeys from relying on local absolute disparity cues to perform the task. The dots were rendered with stereoscopic and perspective cues, and scaled according to the distance so that their screen size only depended on the slant. The baseline dot size was 0.35° isotropic.

### Tilt discrimination task

The monkeys were trained to perform an 8AFC tilt discrimination task (Chang et al., 2020). Each trial began with fixation of a circular target (0.3°) at the center of the screen for 300 ms. A plane then appeared at the center of the screen for 1 s. The target and plane then disappeared, and 8 choice targets appeared at polar angles of 0° to 315° in 45° steps (11° eccentricity). The nearest side of the plane was reported by making a saccade to the corresponding target for a liquid reward. Responses to frontoparallel planes (slant = 0°, tilt undefined) were rewarded with equal probability (12.5%).

### Visually guided saccade task

Each trial began with fixation of a target at the center of the screen for 1.3 s. The fixation target then disappeared and a saccade target appeared at one of the choice target locations. A saccade to the target was made for a liquid reward.

### Experimental protocol

Stimuli were presented in a pseudorandom order within a block design. A block included one completed trial for each of the following: (*i*) planes: (8 tilts x 4 non-zero slants + 8 frontoparallel trials) x 4 distances (N = 160), and (*ii*) saccades: 8 directions x 4 repeats (N = 32). A trial was aborted and data discarded if fixation was broken before the choice or saccade targets appeared, or if the trial was not completed within 500 ms of their appearance. At least five complete blocks were required to include a neuron for analysis.

### Neuronal recordings

Area CIP was identified based on anatomical and functional properties (Rosenberg et al., 2013; Rosenberg and Angelaki, 2014b, a; Elmore et al., 2019). Briefly, the CARET software was used to register the structural MRIs to the F99 atlas (Van Essen et al., 2001) and align the recording grids to estimate electrode trajectories. The area was functionally distinguished from neighboring regions based on 3D orientation selectivity and large receptive fields. Recordings were performed using silicone linear array probes with four or eight tetrodes (NeuroNexus, Inc.). Tetrodes were separated by 300 μm. Electrodes within a tetrode were arranged in a diamond pattern and separated by 25 μm. Neuronal signals were sampled at 30 KHz and stored with eye movement and phototransistor traces sampled at 1 KHz (Scout Processor; Ripple, Inc.). Tetrode-based spike sorting was performed offline using the KlustaKwik (K. Harris) semi-automatic clustering algorithm in *MClust* (MClust-4.0, A.D. Redish et al.) followed by manual refinement using Offline Sorter (Plexon, Inc.). Only well-isolated single neurons verified by at least two authors were included for analysis: 214 neurons from the left hemisphere of Monkey L (26 sessions) and 209 neurons from the right hemisphere of Monkey F (27 sessions).

### Analysis of behavioral data

Tilt discrimination performance was quantified by fitting a von Mises probability density function to the distribution of reported tilt errors (Chang et al., 2020), ΔTilt = reported tilt – presented tilt:

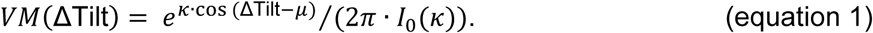

The mean (μ) and concentration (*κ*) parameters describe accuracy and sensitivity, respectively (Seilheimer et al., 2014; Dakin and Rosenberg, 2018). Values of μ closer to 0 indicate greater accuracy. Larger *κ* indicate greater sensitivity. Given the 45° tilt sampling interval, we set *κ* = 18 as the upper bound in the estimation routine (Chang et al., 2020). A modified Bessel function of order 0, *I*_0_(*k*), normalizes the function to have unit area.

### Analyses of neuronal data

#### Visual response latency

To estimate a neuron’s visual response latency, a spike density function (SDF) was created for each trial that a plane was presented by convolving the spike train (1 ms bins, aligned to the stimulus onset) with the function (Schwemmer et al., 2015):

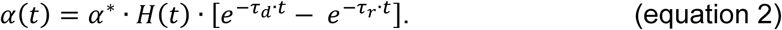

Here, *α*^*^ normalizes *α*(*t*) to have unit area, *H*(*t*) is the Heaviside function, and *τ*_*d*_ = 0.05 and *τ*_*r*_ = 1.05 are time constants. The response latency was defined as the first time point after stimulus onset that the firing rate was significantly different from baseline for at least 30 ms (ANOVA, *p* < 0.05). The baseline firing rate was calculated using the last 150 ms of the fixation periods preceding the stimulus onsets.

#### Choice-related activity

We tested for choice-related activity using the frontoparallel plane data. To combine data across distances, we z-scored all baseline subtracted frontoparallel plane responses at each distance. We then grouped the z-scored firing rates according to the choice. A neuron was classified as having choice activity if the z-scored firing rates significantly depended on the choice (ANOVA, *p* < 0.05). These neurons were used to estimate the time course of choice activity. Average SDFs were calculated for each choice (aligned to the stimulus onset), and labeled relative to the choice that elicited the maximum z-scored firing rate for the neuron: preferred choice, ±45°, ±90°, ±135°, and 180°. The SDFs were averaged across neurons to create eight population-level time courses. The onset of choice activity was defined as the first time point after stimulus onset that the time courses significantly differed (ANOVA, *p* < 0.05). To refine the estimate, we iteratively repeated this process with all neurons, each time calculating the firing rates starting from the previous estimate of the onset of choice activity to the end of the 1 s stimulus presentation (firing rates were first calculated using the full 1 s). This process was repeated until the onset no longer changed.

#### Quantifying orientation selectivity

For each distance and time window, we tested for orientation tuning (ANOVA, *p* < 0.05, with Bonferroni-Holm correction). The preferred tilt and slant was estimated for each significant case by fitting a Bingham function (Rosenberg et al., 2013). Differences in orientation preference were assessed as follows. First, we calculated the principal orientation about which the measured orientation preferences clustered (Fisher et al., 1993). For each distance and time window with significant tuning (N ≤ 8), the surface normal vector *n*_*i*_ = [*x*_*i*_ *y*_*i*_ *z*_*i*_]^*T*^ of the plane with the preferred tilt (*T*_*i*_) and slant (*S*_*i*_) was calculated:

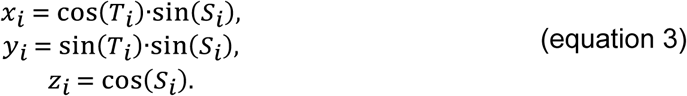

The normal vectors were then arranged in a matrix and the eigenvectors calculated. The principal orientation was defined as the eigenvector with the greatest eigenvalue. Principal orientations were calculated using the SO window only and both (SO+SPC) windows. Second, we rotated the normal vectors such that the principal orientation aligned with the north pole (*n* = [0 0 1]^*T*^):

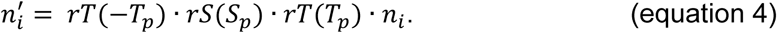

Here, 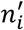 is a rotated normal, *T*_*p*_ and *S*_*p*_ are the tilt and slant of the principal orientation, respectively, and *rT* and *rS* are rotation matrices:

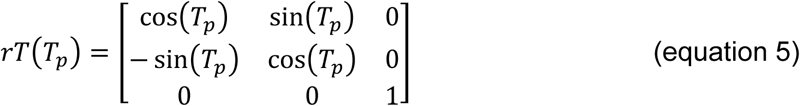

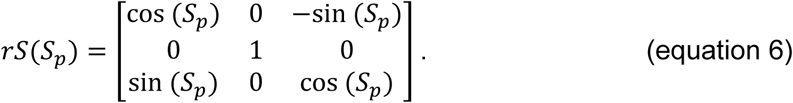

The equatorial projection of the rotated normal vectors is a standardized polar coordinate space that describes deviations in orientation preference.

#### Quantifying tolerance

To quantify how tolerant each neuron’s orientation selectivity was to distance, we fit the 33 × 4 (orientations x distances) matrix of responses with a multiplicatively separable model:

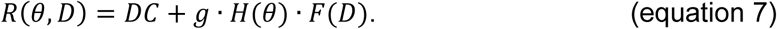

Here, *R*(*θ, D*) is the response to orientation *θ* (tilt and slant) and distance *D, DC* is an offset, *g* is the response gain, *H*(*θ*) is the orientation tuning, and *F*(*D*) is the distance tuning. Fitting was performed using singular value decomposition within a minimization routine to find the *DC, g* (first singular value), and *H* and *F* (first pair of singular vectors) that minimized the Euclidean norm of the error. A tolerance index was defined as the average Pearson correlation between the observed and model orientation tuning curves at each distance. A value of 1 indicates that the shape of the orientation tuning curve was perfectly invariant to distance. Values closer to 0 imply sensitivity to lower-level visual features.

We also tested an additively separable model, *R*(*θ, D*) = *DC* + *H*(*θ*) + *F*(*D*). To fit the model, we constructed a system of equations for the 132 orientation and distance combinations. The closed form solution for the resulting linear regression problem was found by including a regularization parameter that minimized the Euclidian norm of the error (Hastie et al., 2009), giving this model one more free parameter than the multiplicatively separable model. Results with the additive model are not presented because the responses of every neuron in both time windows (846 comparisons) were better described by the multiplicatively separable model.

#### Saccade analysis

A saccade onset was defined as the first time point that the velocity of either eye was ≥ 150°/s. A neuron was classified as having saccade direction selectivity if the baseline subtracted firing rates significantly depended on the saccade direction (ANOVA, *p* < 0.05). These neurons were used to estimate the time course of saccade activity. Average SDFs were calculated for each saccade direction (aligned to the saccade onset), and labeled relative to the saccade direction that elicited the maximum firing rate for the neuron. The SDFs were averaged across neurons to create eight time courses. The onset of saccade activity was defined as the earliest time point before saccade onset that the time courses significantly differed (ANOVA, *p* < 0.05). To refine the estimate, we iteratively repeated this process with all neurons, each time calculating the firing rates from the previous estimate of the onset of saccade activity to the start of the saccade. This process was repeated until the onset no longer changed.

## Acknowledgments

We thank Satchal Postlewaite for help with spike sorting. This work was supported by the Alfred P. Sloan Foundation (FG-2016-6468), Whitehall Foundation (2016-08-18), Greater Milwaukee Foundation (Shaw Scientist Award), and the National Institutes of Health (EY029438). Further support was provided by National Institutes of Health Grant P51OD011106 to the Wisconsin National Primate Research Center, University of Wisconsin–Madison.

